# AlphaFold2 fails to predict protein fold switching

**DOI:** 10.1101/2022.03.08.483439

**Authors:** Devlina Chakravarty, Lauren L. Porter

## Abstract

AlphaFold2 has revolutionized protein structure prediction by leveraging sequence information to rapidly model protein folds with atomic-level accuracy. Nevertheless, previous work has shown that these predictions tend to be inaccurate for structurally heterogeneous proteins. To systematically assess factors that contribute to this inaccuracy, we tested AlphaFold2’s performance on 98 fold-switching proteins, which assume at least two distinct-yet-stable secondary and tertiary structures. Topological similarities were quantified between five predicted and two experimentally determined structures of each fold-switching protein. Overall, 94% of AlphaFold2 predictions captured one experimentally determined conformation but not the other. Despite these biased results, AlphaFold2’s estimated confidences were moderate-to-high for 74% of fold-switching residues, a result that contrasts with overall low confidences for intrinsically disordered proteins, which are also structurally heterogeneous. To investigate factors contributing to this disparity, we quantified sequence variation within the multiple sequence alignments used to generate AlphaFold2’s predictions of fold-switching and intrinsically disordered proteins. Unlike intrinsically disordered regions, whose sequence alignments show low conservation, fold-switching regions had conservation rates statistically similar to canonical single-fold proteins. Furthermore, intrinsically disordered regions had systematically lower prediction confidences than either fold-switching or single-fold proteins, regardless of sequence conservation. AlphaFold2’s high prediction confidences for one fold-switching conformation corroborate previous work showing that machine-learning-based structure predictors fail to capture other fundamental biophysical features of proteins such as their folding pathways. Our results emphasize the need to look at protein structure as an ensemble and suggest that systematic examination of fold-switching sequences may reveal propensities for multiple stable secondary and tertiary structures.

## Introduction

AlphaFold2 has revolutionized protein structure prediction by using sequence information to rapidly model protein folds with atomic-level accuracy^1; 2^. Its predictions are generated by a deep neural network that identifies features of both multiple sequence alignments (MSAs) and experimentally determined protein structures. In other words, AlphaFold2 predictions are generated by a highly sophisticated deep-learning model that excels at recognizing correlations between protein sequence and structure.

AlphaFold2’s approach to protein structure prediction is rooted in a large training set of experimentally determined protein structures. Indeed, without the Protein Data Bank (PDB), a publically available repository of nearly 200,000 protein structures^3^, it would be impossible to predict protein structure through deep learning^4^. Consequently, deep learning-based approaches are likely to miss protein properties that are not apparent from available experimentally determined structures. For example, the conformations accessible to structurally heterogeneous proteins, whose overall secondary and tertiary structures are either unstable or change in response to their environment, cannot be captured in a single protein structure. Thus, it is not surprising that AlphaFold2 often fails to accurately predict the conformations of intrinsically disordered proteins^5-7^, whose structures are highly heterogeneous. AlphaFold2 predictions cover 99% of sequences in the human proteome (https://alphafold.ebi.ac.uk/), but only 58% of residues are modelled with high confidence^5; 7^. Many low-confidence predictions correspond to intrinsically disordered proteins/regions (IDPs/IDRs), often predicted to fold into long filaments^5; 6^. Furthermore, it remains unknown how accurately high-confidence predictions capture the structures of unchararacterized proteins, especially those with few homologs in sequence databases or with sequences dissimilar to proteins represented in the PDB.

Here, we systematically assess whether AlphaFold2 captures the structural heterogeneity of fold-switching proteins. Contrasting IDPs/IDRs, which are natively unstructured, fold-switching proteins have regions of 20 or more contiguous amino acids that either assume distinct stable secondary and tertiary structures under different cellular conditions or populate distinct stable secondary and tertiary structures at equilibrium^8-10^. Thus, the sequences of fold-switching protein regions encode more than one ordered state^9; 11^. As AlphaFold2 maps primary structure (amino acid sequence) to three-dimensional structure, we compared its predictions with experimentally determined protein structures to explore whether it identifies the two stable structures encoded by fold-switching sequences, a single structure, or something else.

## Methods

### Dataset of fold-switching proteins

A set of 98 proteins^9; 12^ that assumed at least two distinct-yet-stable secondary/tertiary structures (folds) was used for the analysis (**Table S1A**). This unique dataset contains protein pairs with extremely high levels of sequence identity (mean 99%/median 100%, **Table S1B**) but regions of 20 or more contiguous residues with different secondary structure configurations, quantified previously^9^ by comparing aligned secondary structure annotations assigned by hydrogen bonding^13^ and backbone torsion angles^14^. Out of the 98, 93 had the alternate fold solved in PDB (**Tables S1A and B**); 91/93 of these proteins have 90% aligned identity or higher; the other 2 are homologs with experimental evidence of fold switching. Similarly, the remaining 5/98 were homologs of fold switchers with only one solved structure in the PDB. These proteins are expected to switch folds because their closely-related homologs do (such as KaiBs from other strains of cyanobacteria with circadian clocks: 4ksoA, 1wwjA, 1r5pA), or were shown to switch folds by methods other than crystallography, *e*.*g*., NMR (Nuclear Magnetic Resonance), CD (circular dichroism), or cryo-EM (cryogenic electron microscopy 1f16A, 3k2sA) ^12^. The sequences of these 5 proteins were used mainly for generating predictions, followed by analyzing prediction scores after modeling (*Assessment of model quality*) and also for conservation scores from the MSA (Multiple Sequence Alignment) generated during prediction (*Conservation scores and rate of evolution using MSA*), but were not used for structural comparisons using Template Model (TM)-scores and Root Mean Square Deviations (RMSD) (*Assessment of model quality*). The structural comparisons between the experimentally determined pairs along with their sequence identities can be seen in Table S1B. Mean/median TM-scores of fold-switching pairs are 0.58/0.63 and RMSDs are 12.6/9.2Å, demonstrating that the experimentally determined structures differ.

### Dataset of intrinsically disordered proteins (IDP) / intrinsically disordered regions (IDR)

A set of 99 proteins was randomly selected from DisProt (https://disprot.org/), a database of experimentally characterized intrinsically disordered proteins, with disordered regions manually curated from the literature^15^. The proteins chosen for the analysis (**Table S2**) had disordered regions ranging from 20-100 residues (to keep their average sizes similar to fold-switching regions); these regions were not located at termini. The set also included the three disordered proteins mentioned in previous work^6^: histone acetyltransferase p300 (Uniprot: Q09472, DisProt: DP00633), CREB-binding protein (Uniprot: Q92793, DisProt: DP02004) and the RNA-binding protein FUS (Uniprot: P35637, DisProt: DP01102).

### AlphaFold2 model generation

The FASTA sequences of fold-switching proteins were extracted from PDB SEQRES records and used as input to the AlphaFold2 structure prediction model^1^, an open-source implementation of the inference pipeline of AlphaFold v2.0 (https://github.com/deepmind/alphafold) maintained on the NIH HPS Biowulf cluster (http://hpc.nih.gov). Ideally, the sequences for the PDB pairs (corresponding to the two folds) would be identical, but for 56 of the pairs the sequences were not identical, usually due to insertions or deletions. Hence we performed modeling for 154 proteins (56*2 sequences with different lengths+37 sequences corresponding exactly to two structures+5 sequences with one solved structure, **Table S3**). The template database contained PDB structures and sequences released until 2021-07-31, which contains both experimentally determined structures from all 93 pairs of fold-switching proteins. The 99 proteins from the DisProt database were modelled with the same parameters as the fold-switching proteins (**Table S2**).

### Assessment of model quality

AlphaFold’s top-scoring models were ranked from 1 to 5 by <u>p</u>er-residue <u>L</u>ocal <u>D</u>istance <u>D</u>ifference <u>T</u>est (pLDDT) scores (a per-residue estimate of the prediction confidence on a scale from 0 – 100), quantified by determining the fraction of predicted Cα distances that lie within their expected intervals. The values correspond to the model’s predicted scores based on the lDDT-Cα metric, a local superposition-free score to assess the atomic displacements of the residues in the model^1^. Models ranked in the top 5 were compared to the original PDB structure using structural alignment as implemented in TM-align^16^, an algorithm for sequence-independent protein structure comparisons. TM-align first generates an optimized residue-to-residue alignment based on secondary structure connections or topology using dynamic programming iterations. An optimal superposition of the two structures was then built on the resulting alignment and TM-score (ranging from 0 to 1) is reported as the measure of overall accuracy of prediction for the models. TM-score > 0.5 implies roughly the same fold^17^, and a higher value indicates a better match. As an alternative measure of structural similarity, we aligned sequences, used the alignment to determined least-square superposition of backbone atoms (C, Cα, O and N), and calculated their RMSD (root-mean-square deviation) using ProFit (Martin, A. C. R., http://www.bioinf.org.uk/software/profit/). These standards of structural similarity were also used by authors of AlphaFold2 to assess the quality of their predictions^1^.

### Ordering conformations

Conformations with higher TM-scores for at least 3/5 AlphaFold2 predictions were designated “Fold1.” Two exceptional cases, 6z4u/5tpn, were designated “Fold1” because they had good/moderate TM-scores (>0.9/0.66) for 2/5 AlphaFold2 predictions, whereas the remaining 3 predictions had moderate/poor TM-scores (< 0.75/0.22) for both folds. This ordering was maintained for the RMSD analysis.

### Clustering in Figure 1

Figure 1 was clustered with k-means clustering, as implemented in the python module scikit-learn^18^. The number of clusters was determined by searching for the first local minimum of the second derivative (*i*.*e*., curvature) with respect to k-means inertia (**Figure S1**). Predictions in Cluster 1 with either TM-score < 0.8 were assigned to Cluster 2; likewise predictions in Cluster 1 with either RMSD < 5Å were also assigned to Cluster 2 (**Figure S4A**). TM-scores of fold-switching regions of proteins from Cluster 1 were determined by excising fold-switching regions from both experimentally determined structures and the 5 AlphaFold models and comparing them with TMalign. Orders of Fold1 and Fold2 were identical as in **Table S1C** except for 5B3Z_A/5BMY_A, all of whose predictions were biased toward Fold2, and thus their ordering was switched. This result is not surprising given that the fold-switching region of this pair was small compared with the whole protein (29/403 residues). Predictions for Fold2 were considered significantly larger if their TM-scores exceeded the TM-scores of Fold1 by at least 0.05, ruling out cases where predictions were equally good for both folds but the TM-score was marginally better for Fold2.

**Figure 1.**
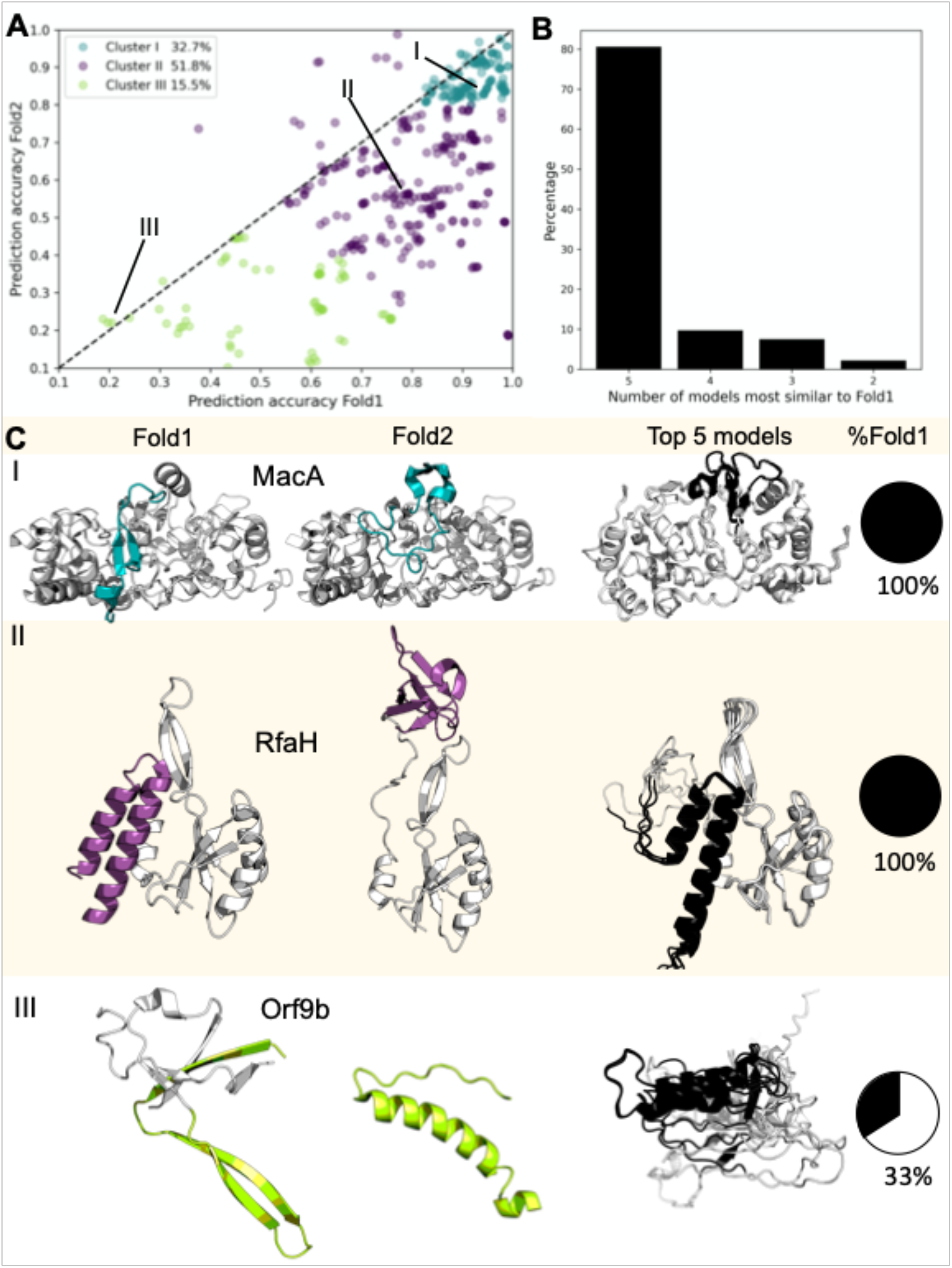
AlphaFold2 fails to predict fold switching. 94% of AlphaFold2 predictions fall below the identity line (dashed line, **A**), indicating bias toward Fold1. Predictions were clustered by the quality of their correspondence with experiment (good predictions with TM ≥ 0.8 for both conformations were colored teal; moderate, purple; poor, green). Furthermore, all 5 of AlphaFold2’s best models corresponded to Fold1 for 81% of fold-switching sequences **(B)**. Experimentally determined structures representing all three clusters are shown in **C** and compared with the top 5 AlphaFold2 predictions for their sequences. Experimentally determined fold switching regions are colored according to their cluster; predicted fold-switching regions are black; single-folding regions are gray. The PDBIDs, chains, length and TM-scores/RMSDs are as follows: (I) 4aanA/4aalA – 341, 0.5/5.2 Å, (II) 2ougC/6c6sD – 162, 3.2/17.5 Å, (III) 6z4uA/7kdtB – 97, 5.0/14.3 Å.

### Fold-switching/single-folding regions

As reported previously^19^, we divided the alignments into fold-switching and single-folding regions, since fold-switching proteins are typically composed of both^9^. Sequences of fold-switching regions along with their lengths are reported in **Table S1A**.

### Conservation scores and rate of evolution using MSA

The multiple sequence alignments (MSAs) generated by AlphaFold2 to predict residue-residue distances and orientations were used to determine evolution rates, using Rate4Site (https://www.tau.ac.il/~itaymay/cp/rate4site.html)^20^. The program requires an MSA file to compute a phylogenetic tree and then calculates the relative conservation score for each column in the MSA. An empirical Bayesian method, which significantly improved the accuracy of conservation score estimates over the Maximum Likelihood method, was used to generate the rates^20^. The scores are represented as grades ranging from conserved-9 to variable-1.

### Statistical tests

The distributions of pLDDT scores and Rate4Site grades in fold-switching regions were compared to the rest of the protein (the single-folding regions) and with a set of 99 IDP/IDRs (**Table S2**). Significance of differences in pLDDT distributions was calculated by employing the two-sample Kolmogorov-Smirnov and Epps-Singleton tests implemented in SciPy^21^. As Rate4Site scores yield a discrete distribution, only the Epps-Singleton test was used.

### Scripts and figures

The scripts used for all analyses were written in Perl and Python; protein figures were generated in PyMOL^22^ and plots in Matplotlib^23^ and seaborn^24^.

## Results

### AlphaFold2 predictions are biased towards one conformation/fold

The five top-scoring AlphaFold2 models of each fold-switching sequence were compared with two distinct, experimentally determined conformations using TM-align, which quantifies similarity of topology and connections between secondary structure elements by TM-score^16^. This metric was used by the authors of AlphaFold2 to assess the accuracies of their models^1^. We systematically termed the more accurately predicted conformation as Fold1 and the less accurately predicted on as Fold2. Generally, when three or more models out of five had higher TM-scores (i.e. were more similar) to one experimentally determined structure, we termed it Fold1 (***Ordering conformations***, Methods); the other experimentally determined structure was termed Fold2. To augment TM-scores we performed RMSD comparions as well (**Figure S4**).

AlphaFold2 models were highly biased toward Fold1. A scatterplot of prediction accuracies, measured by TM-scores (**Figure 1A)**, indicates that nearly 94% of predictions fall below the identity line and are thus more similar to Fold1 than Fold2. The k-means algorithm was used to subdivide the scatterplot into three clusters, corresponding to the first local minimum of k-means inertia curvature with respect to the number of clusters (**Figure 1A, Figure S1, Methods**). To simplify discussion, clusters were ordered by prediction quality rather than size. Cluster I comprised ∼33% of all predictions, which were the most accurate for both conformations (TM-scores ≥ 0.8). Cluster II comprised ∼52% of all predictions and generally paired either one good prediction (TM-score ≥ 0.8) and one moderate prediction (0.6 ≤ TM-score < 0.8) or two moderate predictions. Cluster III comprised the remaining 15.5% of predictions, all of which had at least one poor prediction (TM-score < 0.6).

Structural predictions tended to be conformationally homogeneous. Specifically, all 5 models were most similar to Fold1 in over 80% (75/93) of fold-switching sequences (**Figure 1B**). Additionally, TM-scores of Fold1 and Fold2 were very close (average difference in TM-scores 0.022 ± 0.017) in 13 of the remaining 18 cases, again indicating high levels of structural similarity among AlphaFold2-predicted models. The remaining 5 cases sample both conformations with moderate-to-good accuracy, and representatives are shown in **Figure S2**.

Examples of fold-switching proteins from all three clusters are shown in **Figure 1C**. In Cluster 1, a short region of MacA, a bacterial cytochrome *c* peroxidase, switches folds during reductive activation^25^. AlphaFold2 predicts that its fold-switching region assumes only the oxidized conformation in its 5 best models. Although all models in Cluster 1 had good TM scores (≥0.8) for both conformations, they were more similar to Fold1 than Fold2. Good scores for Fold2 likely result from shorter lengths of fold-switching regions compared to the lengths of the remainder of the protein (**Table S1A**), which had good overall predictions except for the relatively short fold-switching regions. Indeed, TM-scores comparing predicted and experimentally determined fold-switching regions of proteins in Cluster 1 were biased toward Fold1 (**Fig. S3**): only 10% of predictions (15/150) had better TM-scores for Fold2 (**Methods**). Prediction qualities for 14/15 of Fold2-favored predictions were poor for both experimentally determined folds (TM-score < 0.6), demonstrating that AlphaFold2 did not capture either experimentally determined conformation well. Representing Cluster 2, RfaH, a transcription factor that regulates the expression of virulence genes in *E. coli*, has a C-terminal domain (CTD) that completely switches between α-helix and β-sheet folds^26^. AlphaFold2 predicts that its CTD assumes only the autoinhibited α-helix conformation in its top 5 models. In Cluster 3, Orf9b, a protein encoded by Severe Acute Respiratory Syndrome Coronavirus 2 (SARS-CoV-2), assumes both a dimeric β-sheet form and a monomeric α-helical form that binds the mitochondrial host protein Tom7^27^. In 2/5 cases, AlphaFold2 predicts a conformation similar to its β-sheet fold (TM-scores of ∼0.66 for both models). The remaining three models are partially helical but mostly unstructured and do not correspond well to either experimentally determined structure as evidenced by TM-scores ranging from 0.17-0.23 (**Table S1C**). These three poor predictions may result from Orf9b’s shallow alignment of 6 sequences, an inadequate number for generating robust distance restraints^1; 28^. Furthermore, many amyloid-forming proteins, such as alpha-synuclein, amylin, and the αβ42 peptide, also fall into Cluster 3. This result corroborates previous observations that AlphaFold2’s approach is not yet sensitive enough to robustly predict the conformations of fibril-forming proteins^29; 30^.

TM-scores can sometimes mislead, especially when comparing mostly helical segments and/or highly dissimilar structures or sequences. To assess the accuracy of this metric, we used RMSD (root-mean-square deviation) to compare AlphaFold2 models with Fold1 and Fold2, whose ordering was kept identical to TM-score ordering. In other words, for the RMSD calculations, Fold1 and and Fold2 were not reordered by accuracy but rather maintained the same ordering assigned by TM-score (**Methods, *Ordering conformations***). This allows consistent comparison between RMSD and TM-score calculations. It should also be noted that the authors of AlphaFold2 also combined TM-scores and RMSDs to assess the accuracy of their models^1^.

As with TM-scores, AlphaFold2 predictions tended to have better RMSDs from Fold1 than Fold2 (**Figure S4A)**. Specifically, predictions where RMSD was ≤ 5Å for at least one structure were better for Fold1 in 83% of cases. Additionally median/mean RMSDs were significantly more accurate for Fold1 (2.9/5.7Å) than Fold2 (9.6/11.9Å). TM-scores were plotted against sequence identities (calculated on the alignment generated by TM-align) between the protein and the prediction (**Figure S4B**). For ambiguous cases with sequence alignments and TM-scores <0.5, the RMSD values were also large (mostly >10 Å), as seen in the bar plot inset. Hence, structural deviations in these ambiguous cases are corroborated by high RMSD values. Finally, TM-scores and RMSDs of AlphaFold2 models vs. experimentally determined fold switchers were significantly correlated: Pearson R: −0.62, p < 3.3*10^−98^ (assuming normal distribution, **Figure S4C**). Together, these results indicate that AlphaFold2 preferentially predicts one fold-switch conformation over another.

### AlphaFold2 predictions of “ground” and “excited” state conformations

Only a few fold switchers have either been shown to populate two folds simultaneously in solution or populate two distinct crystal forms^10; 26; 31; 32^. Here, we found 7 (**Table S4**), all of which had only 1 conformation predicted by AlphaFold2. More typically, fold-switching proteins assume a more stable “ground” state and a less stable “excited” state^33^. Thus, we classified the remaining 86 protein pairs into “ground” and “excited-state” conformations. We define ground state in three ways: first as isolated protein when the other conformation binds a ligand, second as a preferred conformation suggested by the literature, such as the ground state tetrameric conformation of KaiB^34^, and third as one of two bound conformers (seven cases in **Table S4**). This third definition gives AlphaFold2 the benefit of the doubt when both structures are ligand-bound.

One might argue that if AlphaFold2 captures the ground state, its predictions could reasonably be considered correct. It does so in 76% of cases, but not in the remaining 24%. For example, it predicts that KaiB assumes an “excited” thioredoxin fold^33^ in 5/5 cases. Combining all 93 fold-switch pairs, AlphaFold2 captures the ground state conformation 70% of the time, but misses it in the remaining 30%.

### AlphaFold2 prediction confidences are significantly higher for fold-switching sequences than for intrinsically disordered proteins (IDPs)

As with fold-switching proteins, AlphaFold2 frequently mispredicts the conformations of IDPs^6; 7^. Mispredicted IDP conformations often have low confidence scores^5^, calculated using the <u>p</u>er-residue <u>L</u>ocal <u>D</u>istance <u>D</u>ifference <u>T</u>est (pLDDT), which quantifies the fraction of predicted Cα distances that lie within their expected distance intervals^1^. Higher pLDDT scores indicate good agreement between prediction and expectation; pLDDT scores >90, 70, 50, 0 are considered very high, high, low, and very low, respectively.

AlphaFold2’s prediction confidences were compared for fold-switching, single-folding regions within the fold-switching proteins, and intrinsically disordered regions in IDPs. AlphaFold2 was run on 99 IDP sequences randomly selected from the DisProt database^15^ (**Methods**). The pLDDT scores of predicted IDPs were compared with those of the fold-switching and single-folding protein regions determined previously (**Methods**). **Figure 2** shows that IDPs have lower average pLDDT scores (55 ± 24) than fold-switching (80 ± 20) and single-folding (87 ± 16) sequences. Furthermore, 74%/87% of fold-switching/single-folding residues had good pLDDT scores (≥70), compared with only 30% for IDPs. Finally, the overall distributions of all three sets of sequences were statistically dissimilar (p ∼ 0, Kolmogorov-Smirnov and Epps-Singleton tests). Together, these results demonstrate that, in contrast to IDPs, AlphaFold2 predictions of fold-switching sequences have relatively high confidences, though not quite as high as single-folding protein regions.

**Figure 2.**
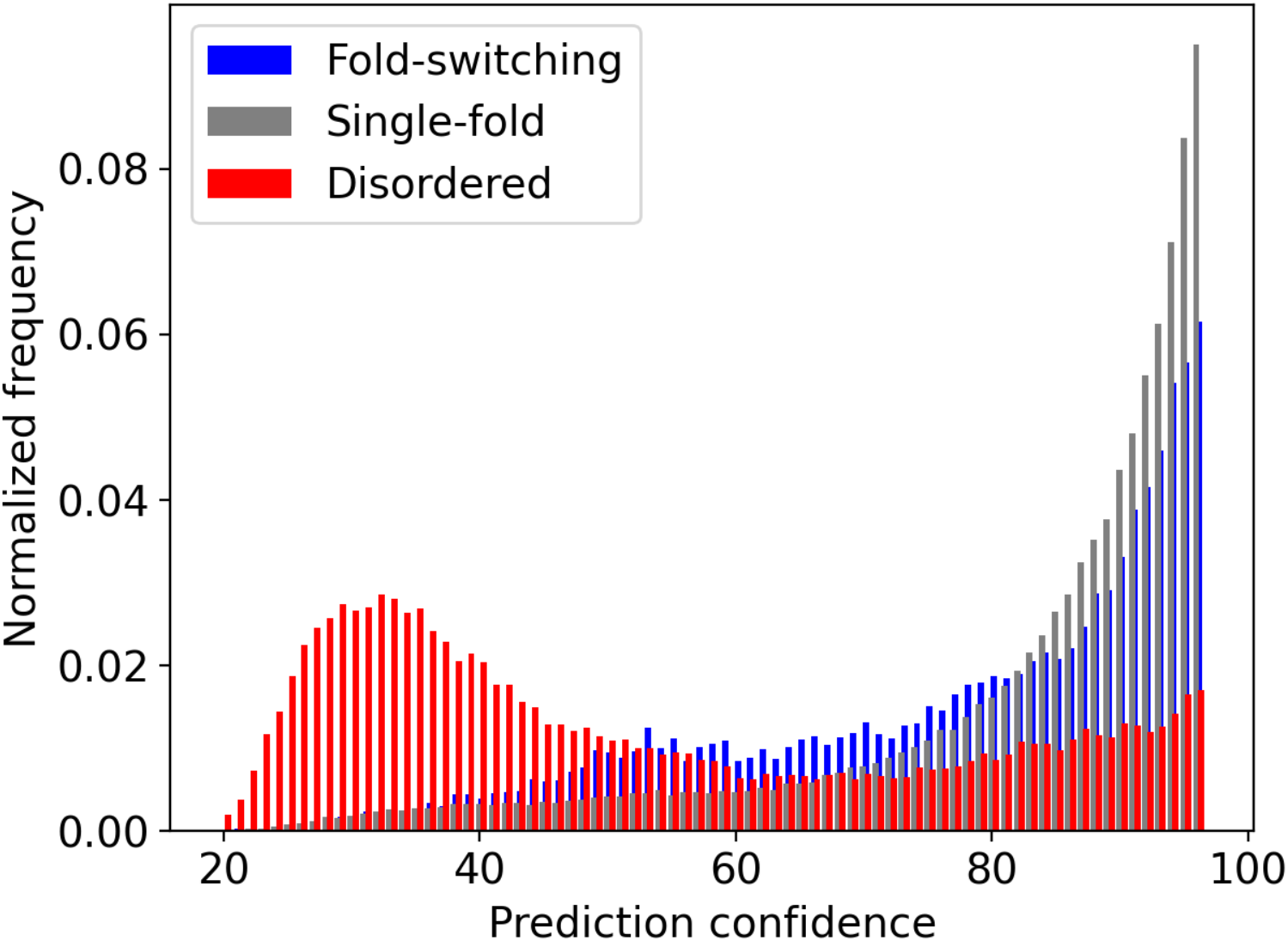
Distributions of AlphaFold2 predictions, measured by pLDDT scores, differ between fold-switching (blue), single-fold (gray), and intrinsically disordered (red) protein sequences. Lower pLDDT scores indicate lower prediction confidences. Thus, AlphaFold2 is generally less confident in its predictions of IDPs than fold-switching or single-folding proteins.

### Fold-switching sequences tend to be more conserved than IDPs

AlphaFold2 does not capture the conformational heterogeneity of IDPs or fold switchers particularly well. While its prediction confidences are generally low for IDPs, they are higher for fold switchers. We investigated whether the low prediction confidences of IDPs resulted from their rapid rates of sequence evolution^35^. This often confounds construction of statistically robust multiple sequence alignments, necessary inputs for generating accurate distance restraints for protein structure prediction^28^. Thus, stronger conservation of fold-switching sequences could be a possible explanation for higher pLDDT scores.

We calculated the conservation scores of the sequences from our set of 98 fold switchers and compared them with 99 IDPs. Specifically, we ran Rate4Site^20^ on multiple sequence alignments generated and used by AlphaFold2 for predictions of fold-switching proteins and IDPs (**Methods**). Distributions of conservation scores, with lower numbers implying less conservation, are shown in **Figure 3**. Conservation scores of single-folding and fold-switching regions were narrowly within the realm of statistical similarity p < 0.074 (Epps-Singleton test, Methods). By contrast, conservation scores of IDP sequences were significantly faster than either fold-switching or single-folding sequences as evidenced by statistically dissimilar distributions with p < 1.1*10^−46^ and 4.4*10^−111^ (Epps-Singleton test, **Methods**) for fold switchers and single folders, respectively.

**Figure 3.**
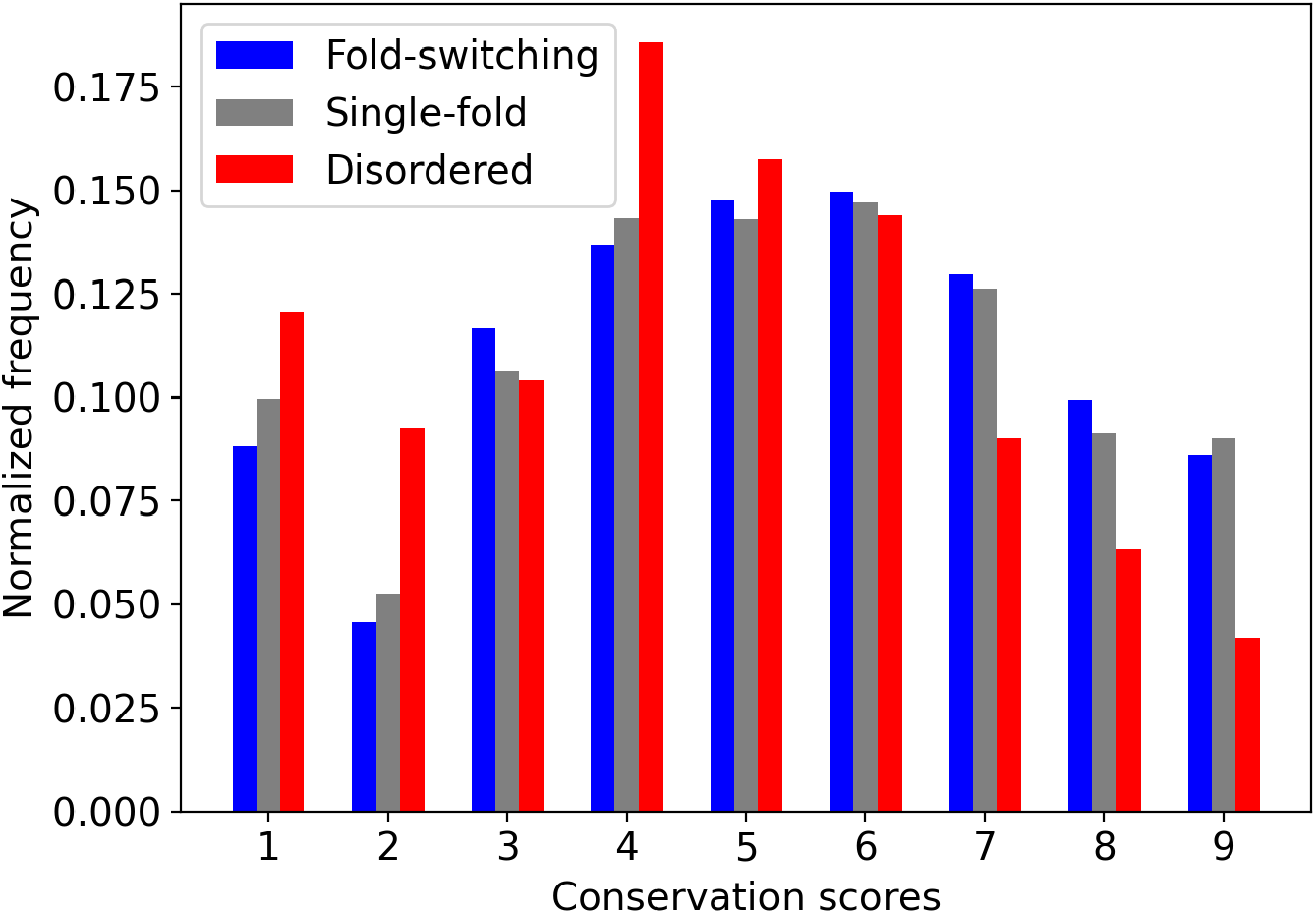
Conservation scores of IDPs, as indicated by Rate4Site grades (1=poorly conserved; 9= well conserved) differ between fold-switching (blue), single-fold (gray), and intrinsically disordered (red) protein sequences. Sequences of fold-switching and single-fold proteins tend to be more conserved than IDP sequences.

These results suggest that AlphaFold2 predictions of fold-switching proteins may have higher confidences because their sequences are more highly conserved than for IDPs.

### Higher prediction confidences suggest that AlphaFold2 searches for one “most probable” conformer

Frequencies of good AlphaFold2 prediction confidences (pLDDT scores ≥ 70) increase with residue conservation for fold-switching, single-folding and disordered proteins (**Figure 4A**). Thus, poorly conserved sequences, specifically conservation scores of 1-3, are associated with lower prediction confidences. At least for IDPs, this is likely explained by poorly conserved sequences having shallower and/or poorly aligned MSAs^1^.

**Figure 4.**
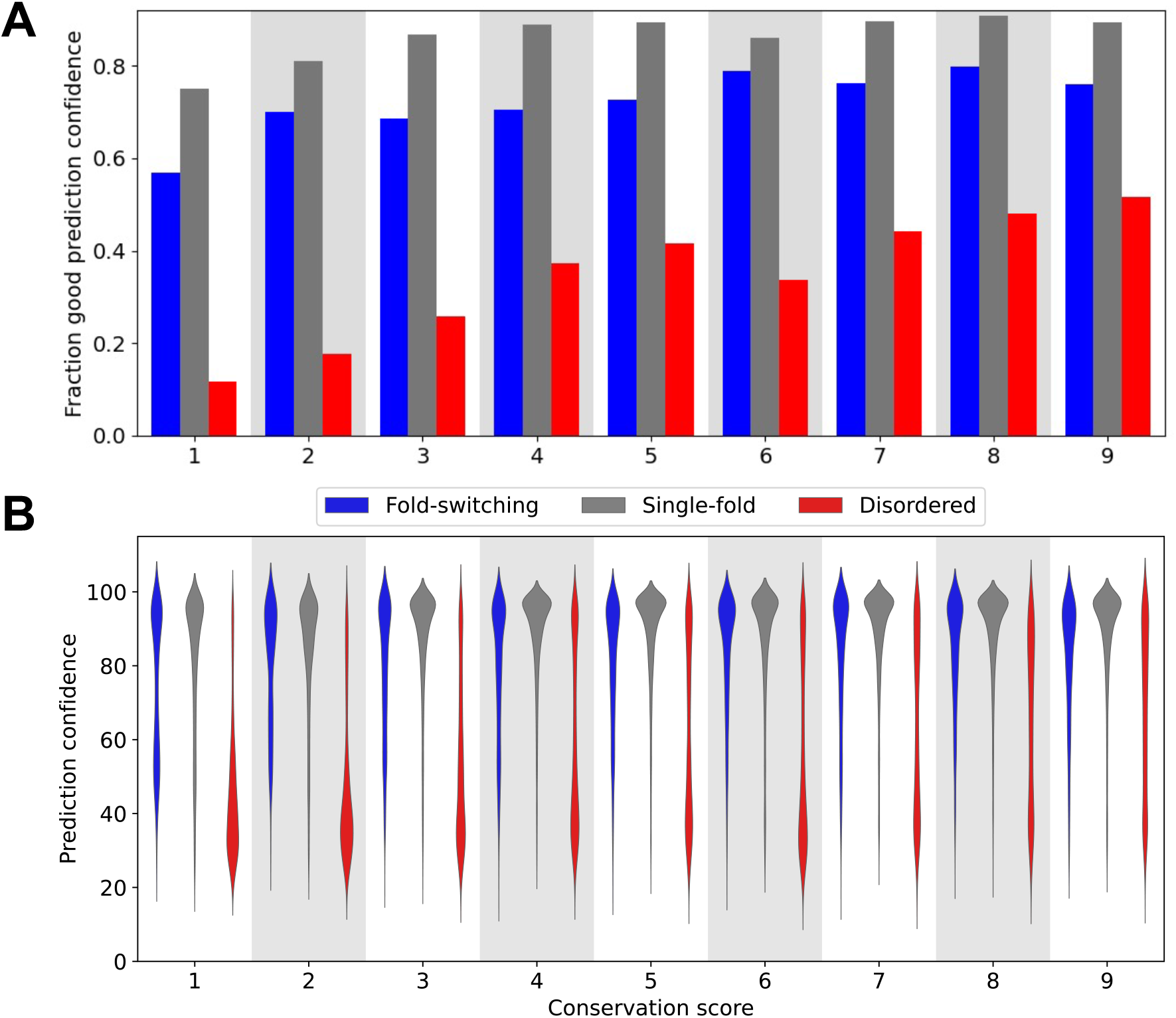
The fraction of AlphaFold2 predictions with pLDDT scores ≥ 70 increases as sequence conservation increases (**A**). Distributions of prediction confidences (quantified by pLDDT scores) are skewed lower for disordered proteins (red) than for single-fold (gray) and fold-switching proteins (blue). (**B**). Wider regions correspond to more populated prediction confidences. In both cases, conservation score was determined using Rate4Site; higher scores correspond to more conserved sequences. Gray/white backgrounds group protein regions with the same conservation score.

Nevertheless, shallow MSAs do not fully explain why AlphaFold2 prediction confidences are higher for fold switchers than for IDPs. Good pLDDT scores level off at around a conservation level of 4 for both fold switchers and single folders, whereas pLDDT scores continue to increase with conservation level for disordered proteins (**Figure 4A**). Furthermore, for all conservation levels, distributions of prediction confidences for disordered residues were skewed systematically lower than corresponding distributions of fold-switching and single-folding residues (**Figure 4B**). Finally, prediction confidences for fold switchers and MSA depth are uncorrelated, as evidenced by a Pearson correlation coefficient of 0.02 (**Figure S5**).

Based on these results, it is likely that AlphaFold2 searches for one “most probable” conformer, instead of an ensemble of possible conformations. This supposition is consistent with 3 additional observations:

1. AlphaFold2 was trained on the PDB, which contains mostly single-fold proteins^9^.
2. AlphaFold2 tends to settle on a protein’s fold early in the prediction process. Specifically, it predicts protein structure using a pairwise representation of amino acid distances in addition to MSAs. These distances are typically determined early in the prediction process and fluctuate minimally with increasing iterations, even for difficult targets^1^.
3. Most structure prediction algorithms assume that proteins assume one stable fold.

## Discussion

AlphaFold2 is a major advance in protein structure prediction^1^, particularly for single-fold proteins^2^. Nevertheless, it is based more on pattern recognition than biophysical principles^4^. Specifically, its deep-learning model is trained on protein sequences and experimentally determined protein structures, neither of which reveal folding mechanisms. Without other fundamental information about protein structure, such as thermodynamics (what balance of forces favor the folded state?) and kinetics (what pathways do proteins traverse between unfolded and folded states?)^4; 36^, deep learning approaches reveal apparent properties of experimentally determined protein structure rather than biophysical pathways^37^. Thus, it is not surprising that its predictions often fail for proteins whose properties are not fully apparent from solved protein structures, such as IDPs^5-7^.

Although AlphaFold2 can be used to predict alternative quaternary structures^38; 39^, here we show that it consistently fails to predict the conformational diversity of fold switchers, proteins that assume multiple secondary structure configurations. Specifically, AlphaFold2 failed to predict fold switching in its top 5 models for 75/93 proteins. Instead, it consistently predicted that fold switchers assume one dominant fold. Since proteins are typically assumed to have one fold, this result is not surprising, especially because AlphaFold2’s training set, the Protein Data Bank, contains relatively few fold-switching proteins^8; 9^. It is notable, however, that AlphaFold2’s predictions miss the ground state of fold switchers 30% of the time. This is further evidence that its predictions are primarily rooted in sophisticated pattern recognition, not protein biophysics^37^.

Unlike IDPs, prediction confidences for fold-switching sequences are relatively high (74% have pLDDT scores > 70 compared with 30% for IDPs). This result, combined with the weak relationship between pLDDT distributions and conservation scores for fold-switching proteins, suggests that AlphaFold2 assumes that stably folded proteins assume one dominant structure. This assumption leads it to miss biologically relevant structural information for some proteins, despite high-confidence predictions. It also raises the question of how much of the full picture AlphaFold2’s full-genome predictions^7^ capture.

The dramatic structural rearrangements of fold-switching proteins regulate biological processes^40^ and are associated with numerous diseases, including COVID-19^27^, cancer^41^, Alzheimer’s^42^, and malaria^31^. Thus, predicting fold-switching proteins is an important problem. While some progress has been made^12; 43; 44^, much work remains to identify features unique to fold-switching proteins. Furthermore, detailed biophysical characterization of fold-switching proteins^45; 46^ is needed. These challenges present an opportunity to improve predictive methods and possibly identify fundamental biophysical principles that are not yet well understood. Such discoveries could help to advance the field of protein structure prediction from sophisticated pattern recognition to methods based fully on protein biophysics.

## Acknowledgements

We thank Loren Looger for critically reading this manuscript. This work utilized resources from the NIH HPS Biowulf cluster (http://hpc.nih.gov), and it was supported by the Intramural Research Program of the National Library of Medicine, National Institutes of Health.

## Supplementary Information

**Fig. S1.**
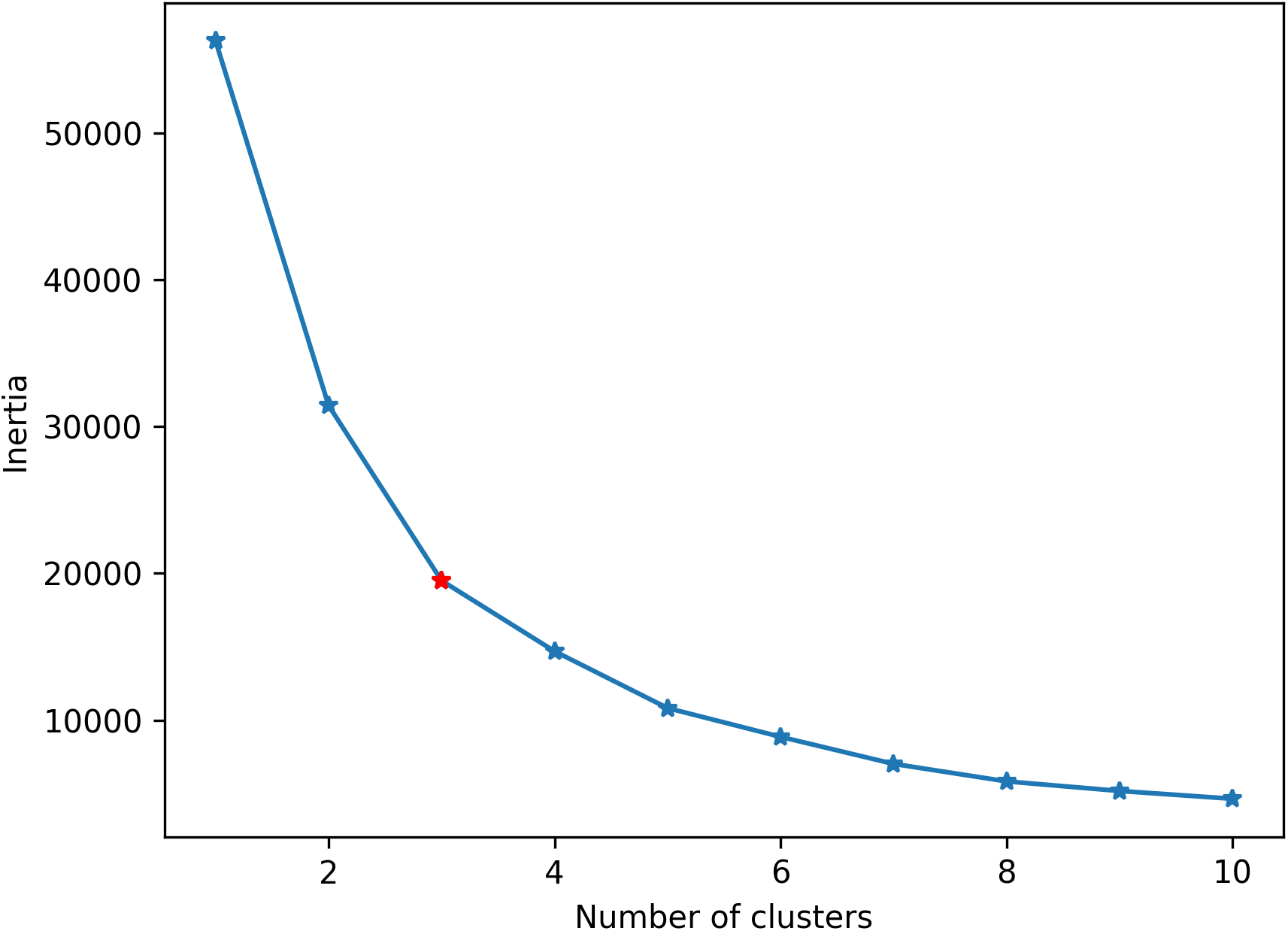
K-means inertia versus number of clusters. The optimal value, determined by finding the minimal number of clusters whose second derivative is less than that of its immediate neighbors, is shown in red.

**Fig. S2.**
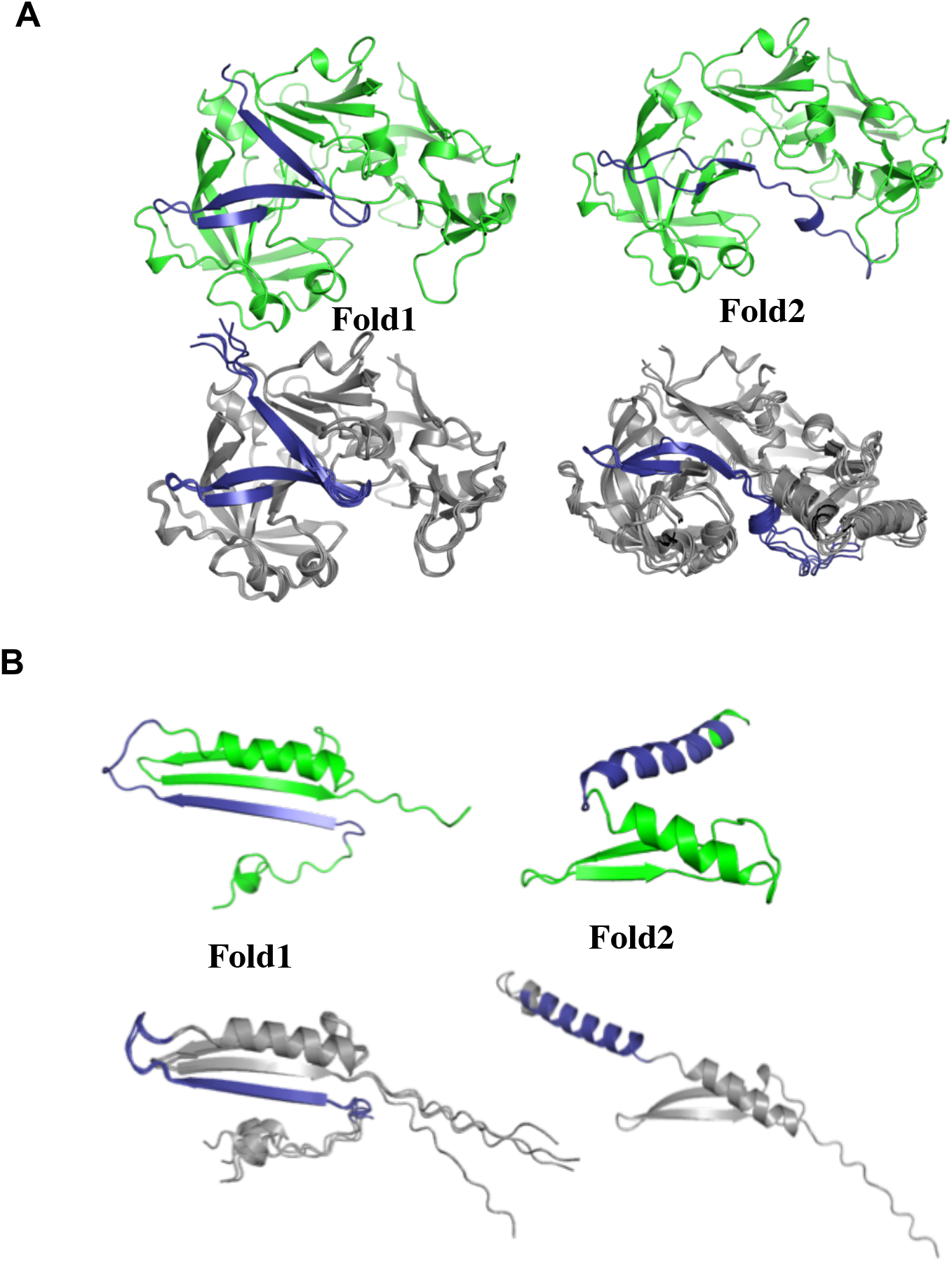
Examples when both folds are present among the predictions. Fold-switching regions are blue in both experimentally-determined and predicted folds (green and gray, respectively). The PDBIDs, chains and TM-scores, RMSDs are as follows: **(A)** Plasmepsin. 1qs8B/1miqB: 0.95/0.93, 1.23/1.46 Å and **(B)** MinE. 2kxoA/3rj9C: 0.77/0.51, 2.4/10.5 Å.

**Fig. S3.**
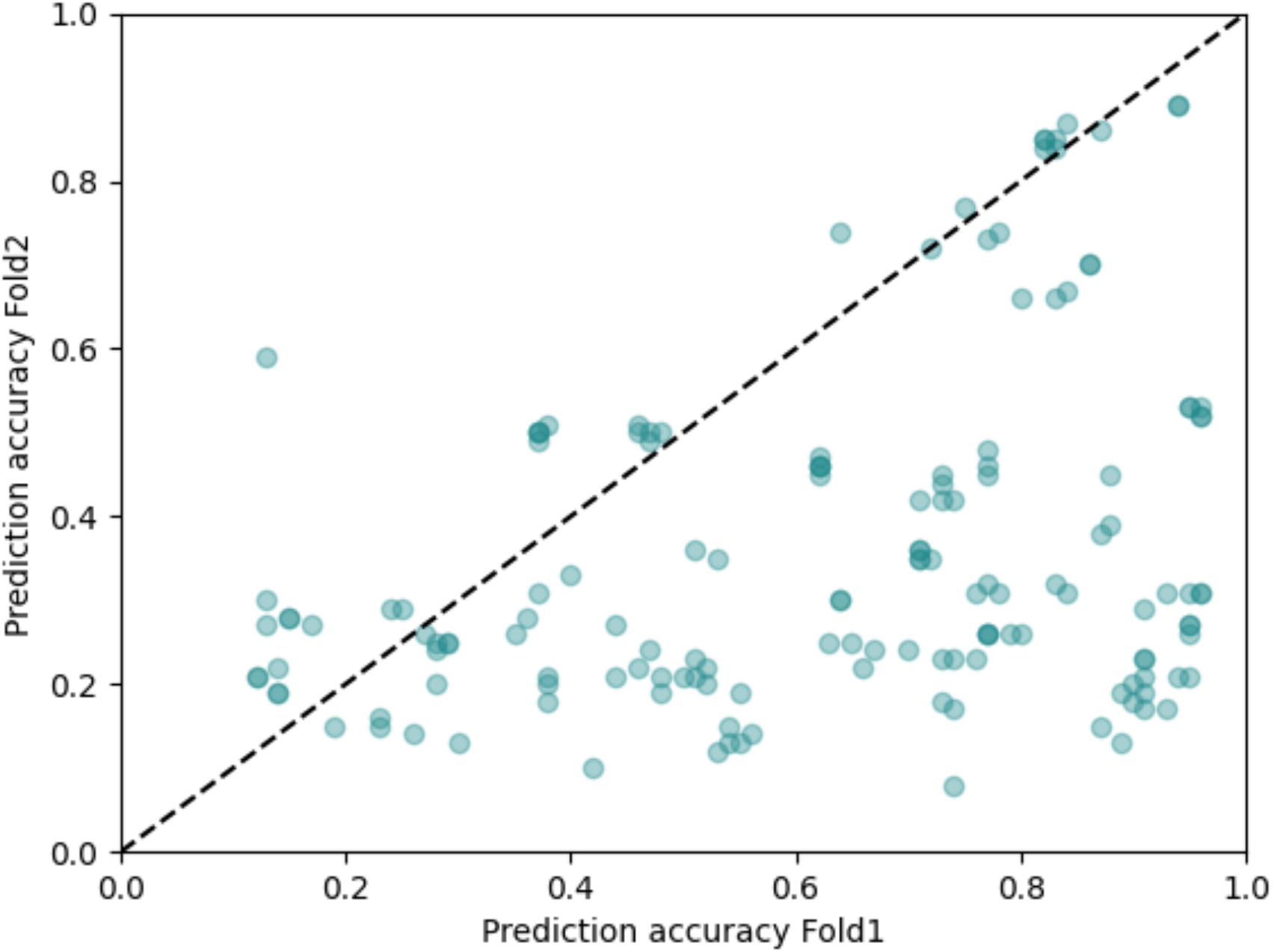
AlphaFold2 predictions of fold-switching regions of proteins from Cluster 1, whose overall folds are predicted well (TM-scores ≥ 0.8, **Figure 1A**), are more accurate for Fold1 than Fold2. Prediction accuracies were quantified using TM-scores, and 41% of predictions were inaccurate (TM-score < 0.6) for both experimentally determined conformations of fold-switching regions.

**Fig. S4.**
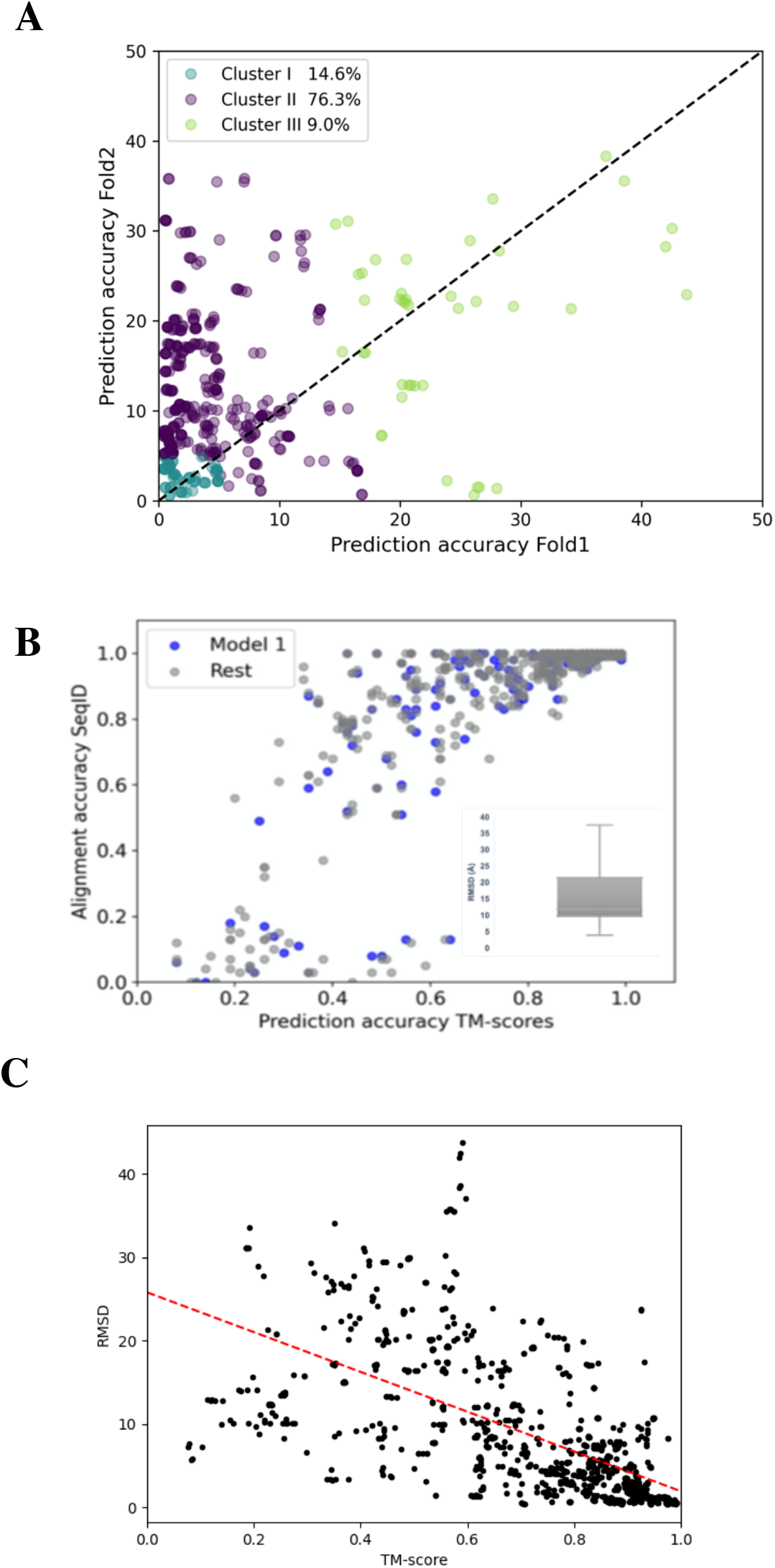
(**A**). Prediction accuracies of fold-switching proteins, assessed by RMSD, are biased toward Fold1. Of the accurately modeled proteins with RMSDs <5Å, 83% are more similar to Fold1 (above the identity line, implying lower RMSD). (**B**) Comparing alignments methods used by TM-align (structural alignment) vs. RMSD (sequence alignment by ProFit followed by structural superposition), for cases having sequence alignment <0.5; RMSD values are presented as box and whiskers plot in inset. (**C**). TM-scores and RMSDs of AlphaFold models and experimentally determined fold-switchers are correlated (r = −0.62, calculated from differences in datapoints from red linear fit). The negative slope is expected since high TM-scores and low RMSDs imply high prediction accuracies.

**Figure S5.**
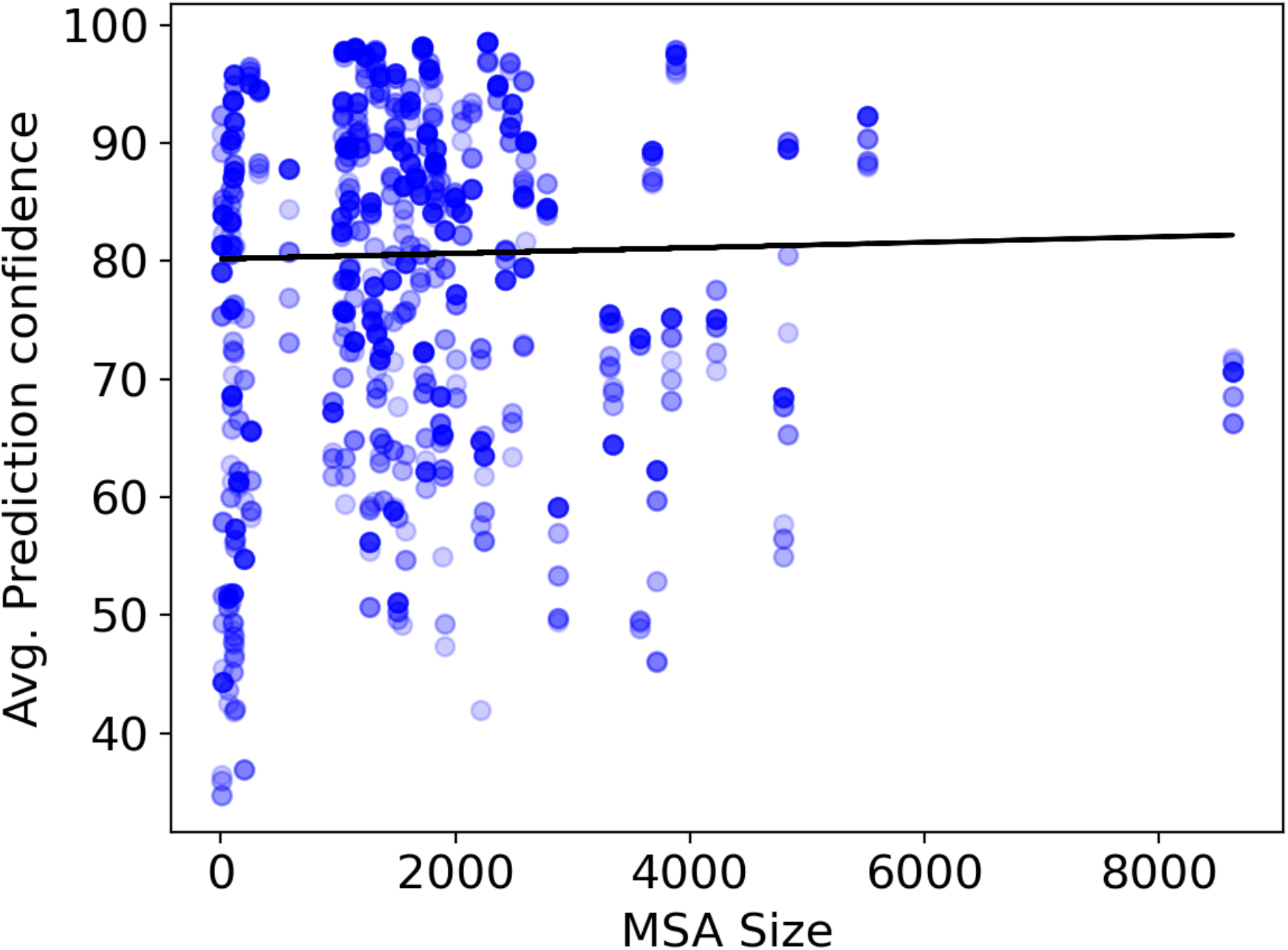
AlphaFold2 prediction confidences (pLDDT scores) are uncorrelated with MSA size (number of sequences in multiple sequence alignment), as evidenced by a Pearson correlation of 0.02 (calculated from differences in datapoints from black linear fit).

**Table S1. (**A) List of pairs (PDBIDs), lengths, sequence of the fold-switching region, methods of structure determination, and resolution (if applicable). Forpairs not having the second fold solved in PDB, only the first PDB is reported. (**B**) RMSD, TM-scores for the whole protein and only fold-switching fragment of experimentally determined fold-switching conformations. Sequence identities are also included. **(C)** List of fold-switching protein pairs (PDBID and chain) used for the analysis; first column corresponds to Fold1 and second to Fold2, followed by TM-scores of the the top 5 predicted models. Tables attached separately.

**Table S2**. Disordered proteins analyzed. Columns correspond to Disprot IDs, Uniprot IDs, Disordered region, and full Uniprot sequence. Attached separately.

**Table S3**. List of pairs in which sequences are not identical in length and sequence, where structure prediction was performed using both sequences for the pair. Attached separately.

**Table S4**. Ground and excited states for all fold-switch pairs. Pairs likely to sample both folds at equilibrium are bold. ** denotes excited state. AlphaFold2 predictions of pairs with 2 excited states were considered to capture the ground state.

